## Supplementary information for "Age-related demethylation of the TDP-43 autoregulatory region in the human motor cortex"

### Supp. Fig. 1

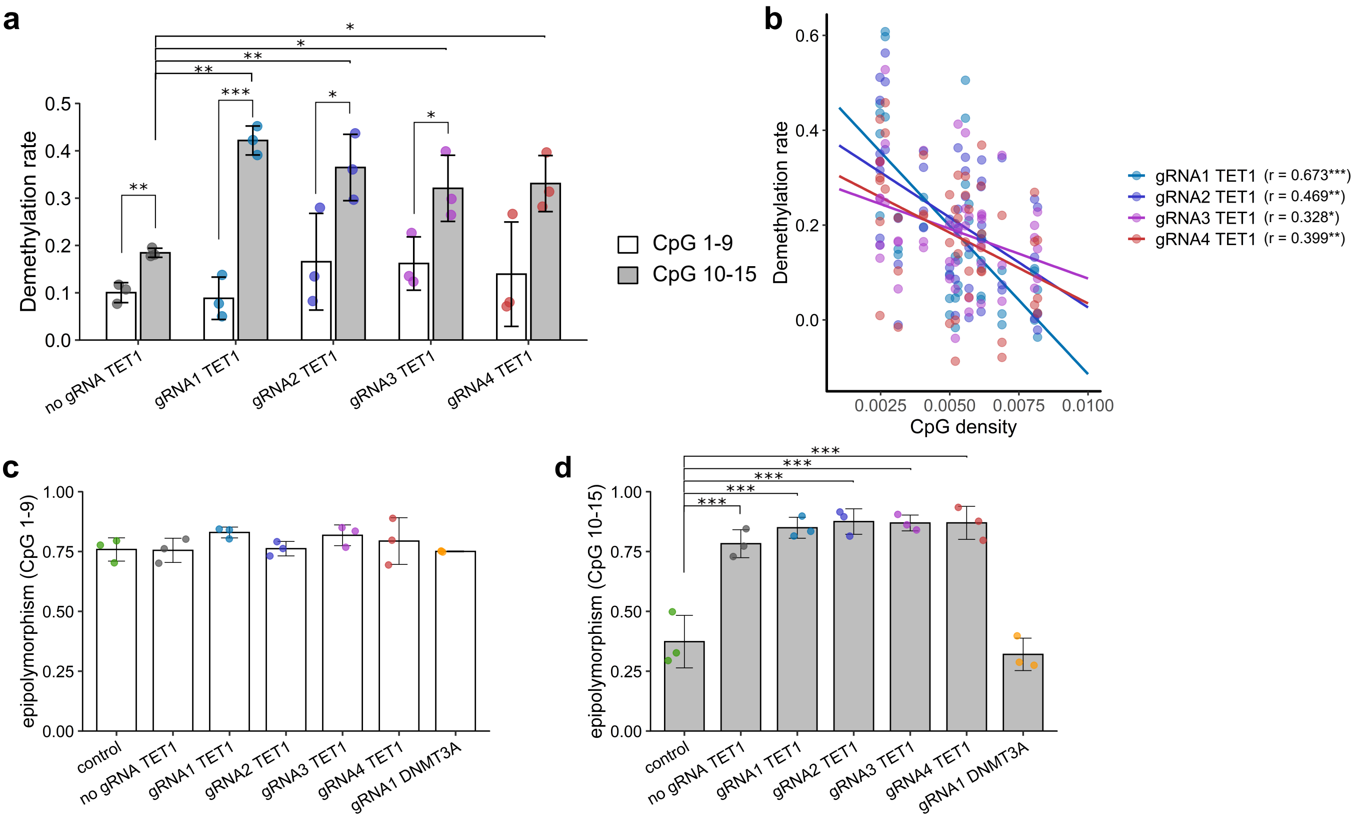


**Supplementary Fig. 1 Demethylation with the TET1-TARDBP-target vector at 3'UTR CpGs 1–15.** (a) Demethylation rate in the controls at CpGs 1–9 and CpGs 10–15 (mean±SD, n=3). Student's t-test; comparison of CpGs 1–9 and CpGs 10–15. Dunnett's test; each guide RNA for no guide RNA. (b) Correlation between the demethylation rate and the CpG density at each CpG site. Pearson's correlation test. (c)(d) Epipolymorphism scores of CpGs 1–9 and CpGs 10–15. Dunnett's test relative to the controls. *p<0.05, **p<0.01, ****p<0.001.

### Supp. Fig. 2

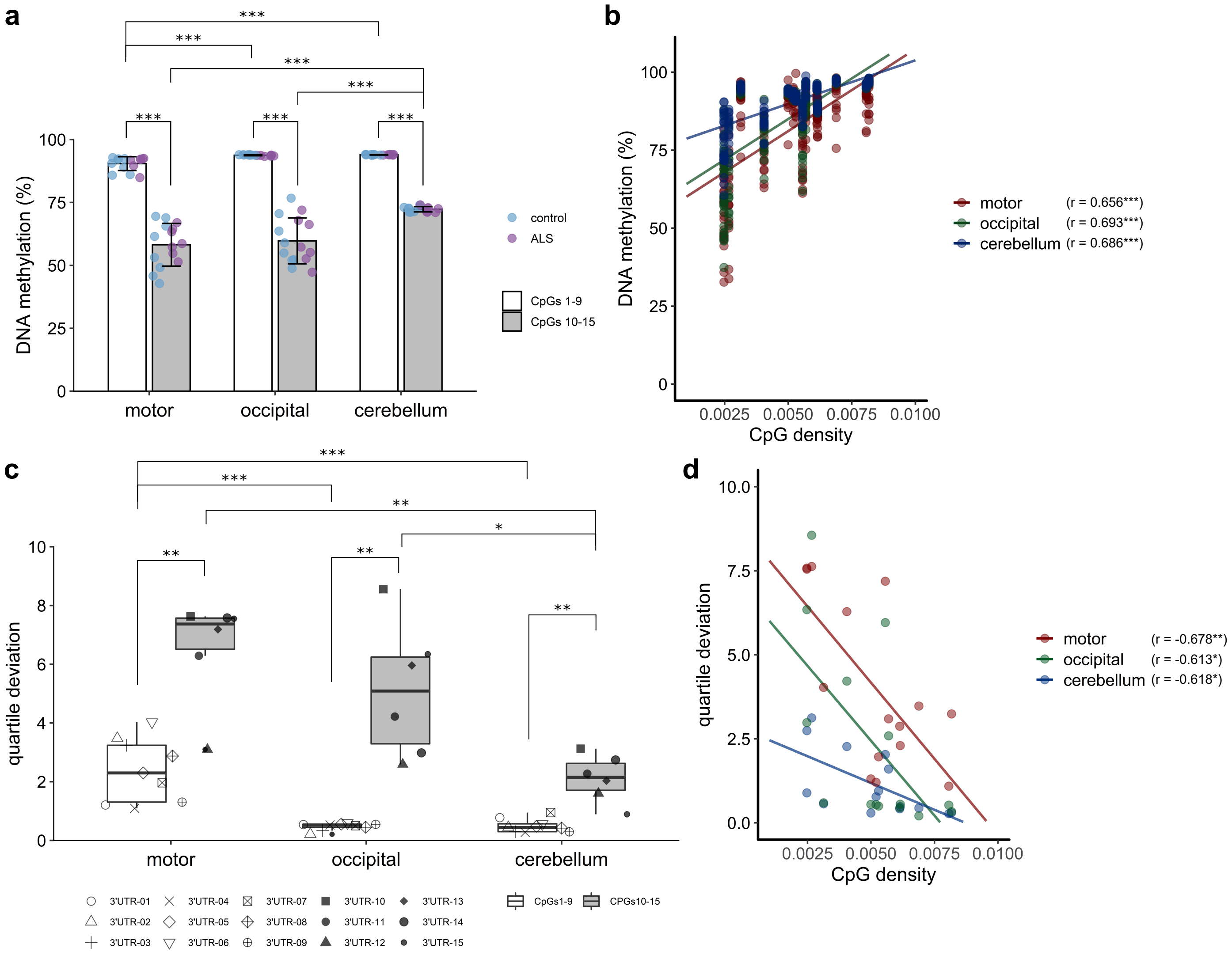


**Supplementary Fig. 2 DNA methylation of the TARDBP gene in the human brain.** (a) Mean DNA methylation percentages of 3'UTR CpGs 1–9 and 3'UTR CpGs 10–15 (mean±SD). (b) Correlation between the percent DNA methylation and the CpG density at each CpG site. (c) Quartile deviation of the percentage of DNA methylation at each CpG site. Box plots of 3'UTR CpGs 1–9 and 3'UTR CpGs 10–15 in each brain region are shown; Welch's t-test was used for the comparisons between 3'UTR CpGs 1–9 and 3'UTR CpGs 10–15; Bonferroni's multiple comparison test was used for the comparisons across the brain regions. (d) Correlation between the quartile deviation of the percentage of DNA methylation and the CpG density at each CpG site. Pearson's correlation test. Seven ALS patients and 8 controls. *p<0.05, **p<0.01, ****p<0.001.

### Supp. Fig. 3

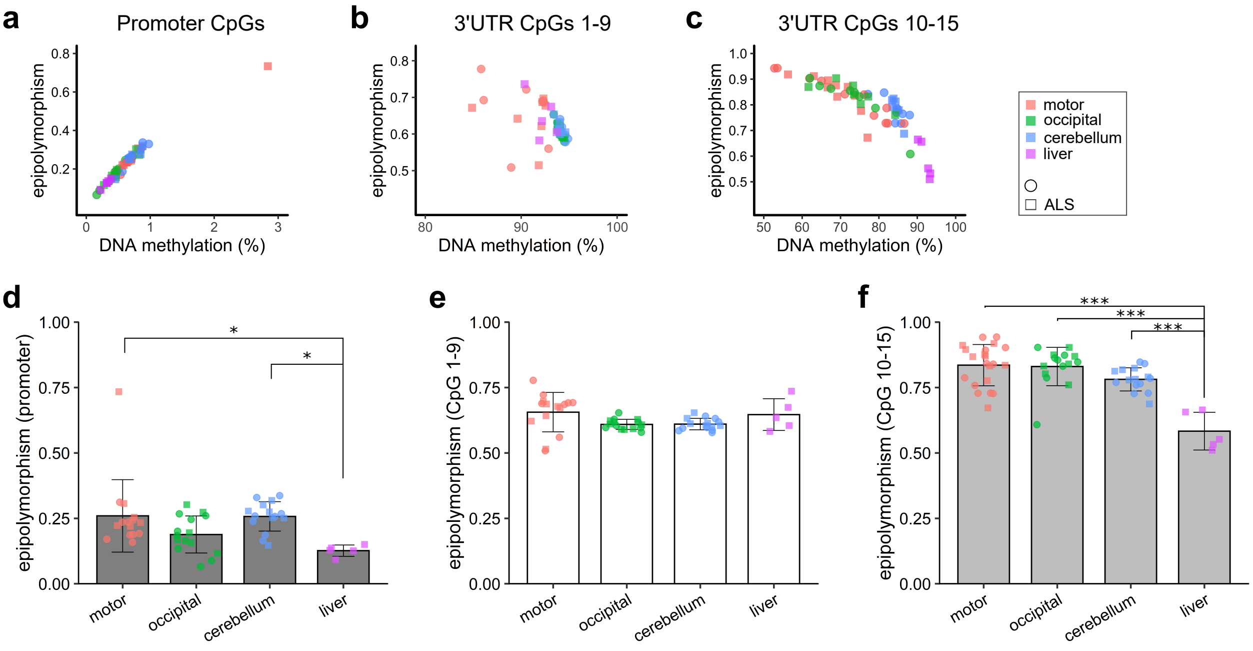


**Supplementary Fig. 3 Epipolymorphism scores of each brain region and liver tissues.** (a)-(c) Average DNA methylation percentages and epipolymorphism scores of promoter regions (a), 3'UTR CpGs 1–9 (b), and 3'UTR CpGs 10–15 (c). Epipolymorphism scores of promoter regions (d), 3'UTR CpGs 1–9 (e), and 3'UTR CpGs 10–15 (f) (mean±SD; Bonferroni's multiple comparison test). Ten ALS patients and 11 controls were included in the analysis of 3'UTR CpGs 10–15 in the motor cortex, five ALS patients were included in the analysis of the liver, and seven ALS patients and eight controls were included in the other analyses. ***p<0.001.

### Supp. Fig. 4

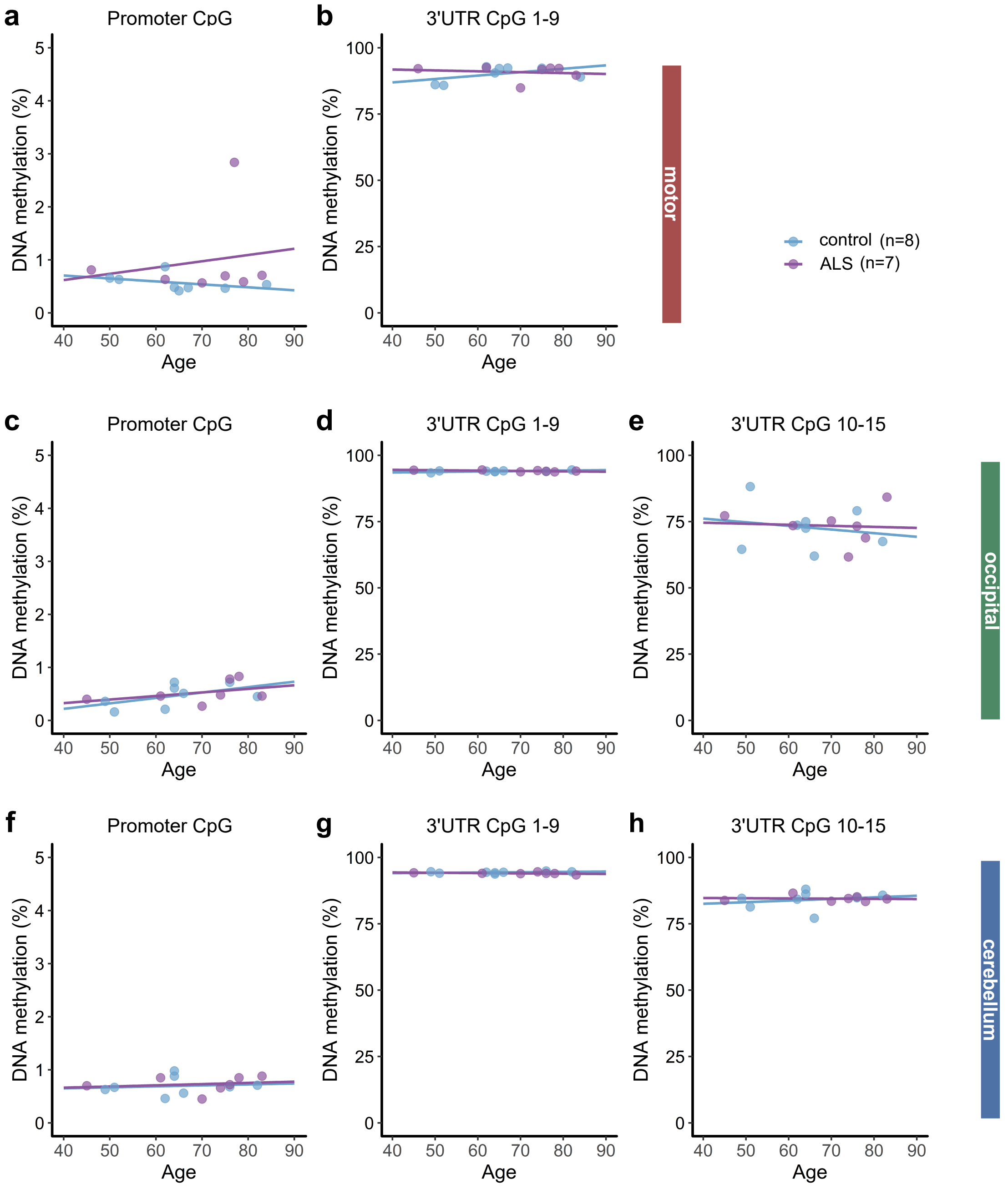


**Supplementary Fig. 4 Scatter plot showing the DNA methylation of the TARDBP gene in the motor cortex (except for 3'UTR CpGs 10-15), occipital cortex and cerebellum according to age at autopsy.** (a-h) Age at autopsy and mean DNA methylation percentages among promoter regions (a, c, f), 3'UTR CpGs 1–9 (b, d, g), and 3'UTR CpGs 10–15 (e, h). (a, b) Motor cortex, (c-e) occipital cortex, (f-h) cerebellum.

### Supp. Fig. 5

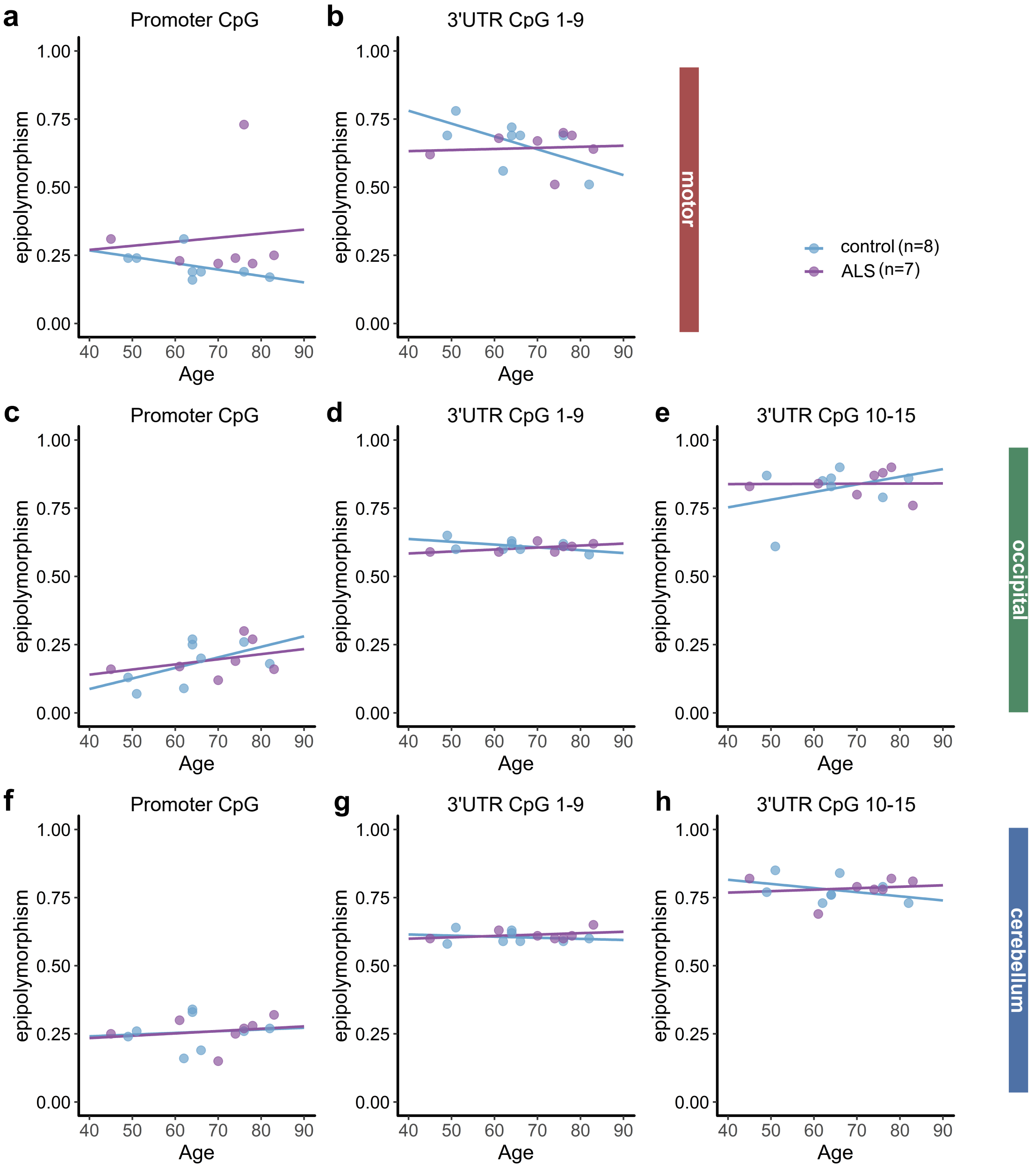


**Supplementary Fig. 5 Scatter plot showing the epipolymorphism score of the TARDBP gene according to age at autopsy.** (a-h) Age at autopsy and epipolymorphism scores of promoter regions (a, c, f), 3'UTR CpGs 1–9 (b, d, g), and 3'UTR CpGs 10–15 (e, h). (a-b) Motor cortex, (c-e) occipital cortex, and (f-h) cerebellum.

#### Supplementary Table 1. Clinical information of the postmortem cases

| Case | Age | Sex | PMI (h:m) | Disease | Age of onset* | Disease duration* (month) | Initial symptom* | DNA methylation analysis | | | | RNA analysis (RIN) | Western blotting |
|---|---|---|---|---|---|---|---|---|---|---|---|---|---|
| | | | | | | | | motor cortex | occipital cortex | cerebellum | liver | motor cortex | motor cortex |
| ALS1 | 78 | M | 2:50 | ALS | 77 | 7 | respiratory dysfunction | + | + | + | + | 9.4 | - |
| ALS2 | 61 | F | 4:00 | ALS | 57 | 46 | upper limb | + | + | + | + | 8.0 | - |
| ALS3 | 76 | M | 1:30 | ALS | 74 | 20 | bulbar | + | + | + | + | 7.7 | - |
| ALS4 | 45 | F | 6:30 | ALS | 44 | 17 | upper limb + bulbar | + | + | + | + | 6.2 | - |
| ALS5 | 83 | M | 2:30 | ALS | 82 | 15 | upper limb | + | + | + | + | 8.7 | - |
| ALS6 | 74 | F | 3:00 | ALS | 71 | 33 | upper limb | + | + | + | - | - | + |
| ALS7 | 70 | F | 1:50 | ALS | 68 | 18 | bulbar | + | + | + | - | 8.9 | + |
| ALS8 | 64 | M | 3:00 | ALS | 62 | 17 | upper limb | + | - | - | - | - | + |
| ALS9 | 84 | F | 5:15 | ALS | 82 | 21 | bulbar | + | - | - | - | 6.7 | + |
| ALS10 | 59 | M | 3:00 | ALS | 51 | 54 | upper limb | + | - | - | - | 8.3 | + |
| control1 | 66 | M | 3:30 | pellagra | - | - | - | + | + | + | - | - | + |
| control2 | 76 | M | 3:30 | gastrointestinal bleeding | - | - | - | + | + | + | - | 7.2 | + |
| control3 | 64 | M | 2:00 | renal failure | - | - | - | + | + | + | - | 8.4 | + |
| control4 | 64 | F | 2:00 | polymyositis | - | - | - | + | + | + | - | 9.1 | + |
| control5 | 51 | M | 4:00 | myopathy | - | - | - | + | + | + | - | 7.3 | + |
| control6 | 82 | F | 4:30 | myasthenia gravis | - | - | - | + | + | + | - | 7.7 | + |
| control7 | 49 | F | 2:00 | POEMS syndrome | - | - | - | + | + | + | - | 9.0 | + |
| control8 | 62 | M | 9:00 | intermittent porphyria | - | - | - | + | + | + | - | - | + |
| control9 | 71 | M | - | renal failure | - | - | - | + | - | - | - | - | + |
| control10 | 54 | M | 1:15 | heart failure | - | - | - | + | - | - | - | 8.0 | + |
| control11 | 68 | M | 3:00 | lung cancer | - | - | - | + | - | - | - | 7.7 | + |

PMI (h:m): post mortem interval (hours: minutes), RIN: RNA integrity number
*Applicable to ALS cases only

#### Supplementary Table 2. Guide RNA target, PCR primer, and probe sequences

| Name | Sequence (5' to 3') |
|---|---|
| guide RNA target sequence *TARDBP* 3'-UTR-1 (except PAM) | ATTCGTCATCACGCATCAC |
| guide RNA target sequence *TARDBP* 3'-UTR-2 (except PAM) | CATCTCATTTCAAATGTTTA |
| guide RNA target sequence *TARDBP* 3'-UTR-3 (except PAM) | CTCCTGTAATATTTTATCCC |
| guide RNA target sequence *TARDBP* 3'-UTR-4 (except PAM) | AATTCTTTGCATGTTCAAAA |
| *TARDBP* promoter primer set 1st F (bisulfite sequencing) | GTTGTTTTTTAGAAAAGGGGTTTAG |
| *TARDBP* promoter primer set 1st R (bisulfite sequencing) | AACTATATAAAAACTAACCTCCCCC |
| *TARDBP* promoter primer set 2nd F (bisulfite sequencing) | ACACTCTTTCCCTACACGACGCTCTTCCGATCTGGTTTTATTTTGTTTTTTAGGTGGATT |
| *TARDBP* promoter primer set 2nd R (bisulfite sequencing) | GTGACTGGAGTTCAGACGTGTGCTCTTCCGATCTGGAACTATATAAAAACTAACCTCCCCC |
| *TARDBP* 3'-UTR primer setA 1st F (bisulfite sequencing) | GTATTTTATTGAAAGTAGTGTTGTAAA |
| *TARDBP* 3'-UTR primer setA 1st R (bisulfite sequencing) | CACCATACAACATTCACAACAATTA |
| *TARDBP* 3'-UTR primer setA 2nd F (bisulfite sequencing) | ACACTCTTTCCCTACACGACGCTCTTCCGATCTGGTATAGGAATATTGTTTATATGTTTTTTT |
| *TARDBP* 3'-UTR primer setA 2nd R (bisulfite sequencing) | GTGACTGGAGTTCAGACGTGTGCTCTTCCGATCTGGCTTCTAATTCCATATCACAACCTTA |
| *TARDBP* 3'-UTR primer setB 1st F (bisulfite sequencing) | GTTGTGATATGGAATTAGAAGGTT |
| TARDBP 3'-UTR primer setB 1st R (bisulfite sequencing) | CACCATACAACATTCACAACAATTA (same as exon6-1 bisulfite 1st R ) |
| *TARDBP* 3'-UTR primer setB 2nd F (bisulfite sequencing) | ACACTCTTTCCCTACACGACGCTCTTCCGATCTGGGTGTGGGATTGATGGTGGT |
| *TARDBP* 3'-UTR primer setB 2nd R (bisulfite sequencing) | GTGACTGGAGTTCAGACGTGTGCTCTTCCGATCTGGATTTCACTTAACACATTCATCTAAT |
| *TARDBP* F1 (RT-PCR) | GCGCTGTACAGAGGACATGA |
| *TARDBP* R1 (RT-PCR) | GCCTGTGATGCGTGATGA |
| *TARDBP* R2 (RT-PCR) | AGTTCCATCTCAAAAGGGTC |
| *TARDBP* unspliced F (qPCR) | TGTCACAGTGTTTGGTTCTTTTG |
| *TARDBP* unspliced R (qPCR) | AGCGGATAAAAATGGGACAC |
| *RPLP1* F and R (qPCR, droplet digital PCR) | Purchased (Takara bio) primer set ID: HA067802 |
| *RPLP2* F and R (qPCR) | Purchased (Takara bio) primer set ID: HA067804 |
| *TARDBP* mRNA F (droplet digital PCR) | CTGCGGGAGTTCTTCTCTCA |
| *TARDBP* mRNA R (droplet digital PCR) | CGCAATCTGATCATCTGCAA |
| *TARDBP* mRNA probe (droplet digital PCR) | CCCAAGCCATTCAGGGCCTTTG |
| *RPLP1* probe (droplet digital PCR) | ACTGCTGCTGCTCCAGCTGA |
